## Supplementary Figures for "A single-cell transcriptomic atlas reveals resident dendritic-like cells in the zebrafish brain parenchyma"

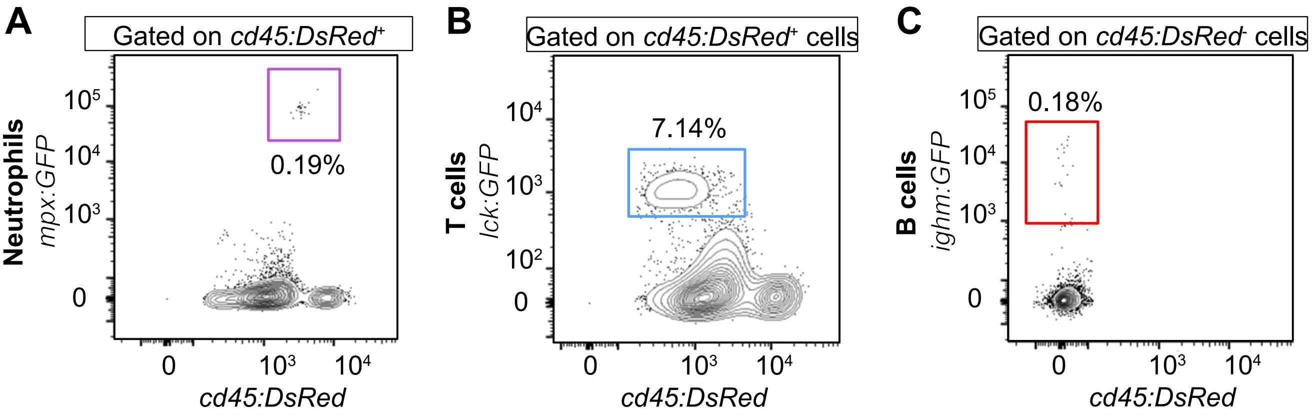


**Figure 1 – supplement 1. Flow cytometry analyses of brain leukocytes using blood lineage-specific GFP reporter lines. A.** Proportion of neutrophils (purple gate), identified using the *Tg(mpx:GFP; cd45:DsRed)* double transgenic line. **B.** Proportion of T/NK cells (blue gate), as determined using Tg(*lck*:GFP; *cd45:DsRed*) fish. **C.** B lymphocytes (red gate), analyzed using *Tg(ighm:GFP; cd45:DsRed)* animals. Note that, as previously shown, the *cd45:DsRed* transgene is not expressed in *ighm:GFP*$^{+}$ B cells. Percentages of each population refer to a single individual and are relative to the total *cd45:DsRed*$^{+}$ population (mean ± SEM of 4 fish: see text).

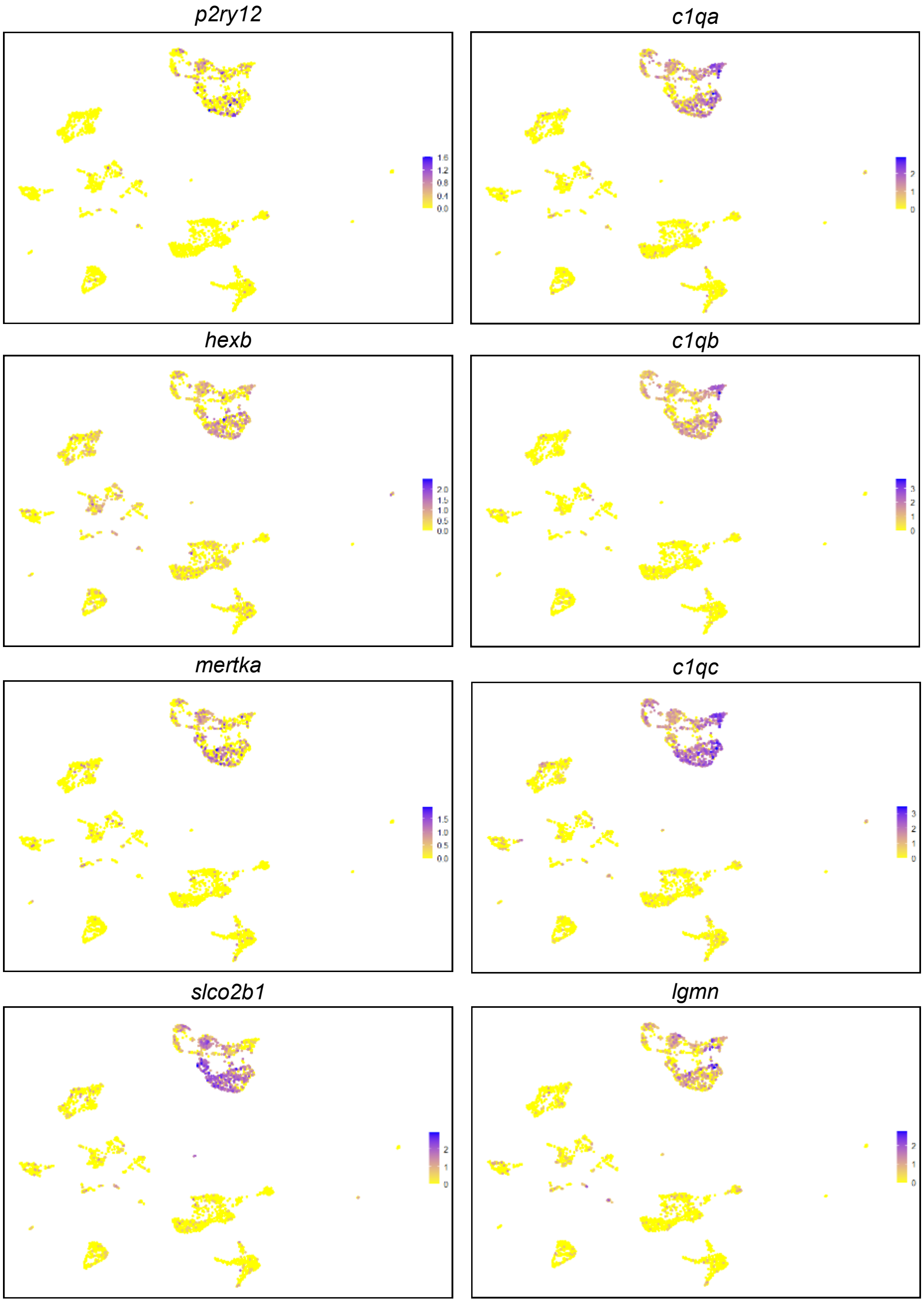


**Figure 4 – figure supplement 1. Canonical microglial genes conserved between zebrafish and mammals.** UMAP plots depicting the expression pattern of zebrafish orthologs of well-known mammalian microglia signature genes (with mammalian orthologs indicated in parenthesis). From top to bottom: *p2ry12* (P2RY12) (Butovsky et al. 2013, Gerrits et al. 2019, Jurga, Paleczna, and Kuter 2020, Van Hove et al. 2019), *hexb* (HEXB) (Butovsky and Weiner 2018, Gerrits et al. 2019, Jurga, Paleczna, and Kuter 2020, Van Hove et al. 2019), *mertka* (MERTK) (Butovsky and Weiner 2018, Gerrits et al. 2019,Jurga, Paleczna, and Kuter 2020, Van Hove et al. 2019), *slco2b1* (SLCO2B1)(Gerrits et al. 2019, Van Hove et al. 2019), *c1qa, c1qb, c1qc* (C1Q A-C)(Gerrits et al. 2019, Butovsky and Weiner 2018, Van Hove et al. 2019), *lgmn* (LGMN)(Gerrits et al. 2019, Butovsky et al. 2013, Van Hove et al. 2019). Color scale (gradual from yellow to purple) indicates the expression level for each gene (normalized counts in log1p).

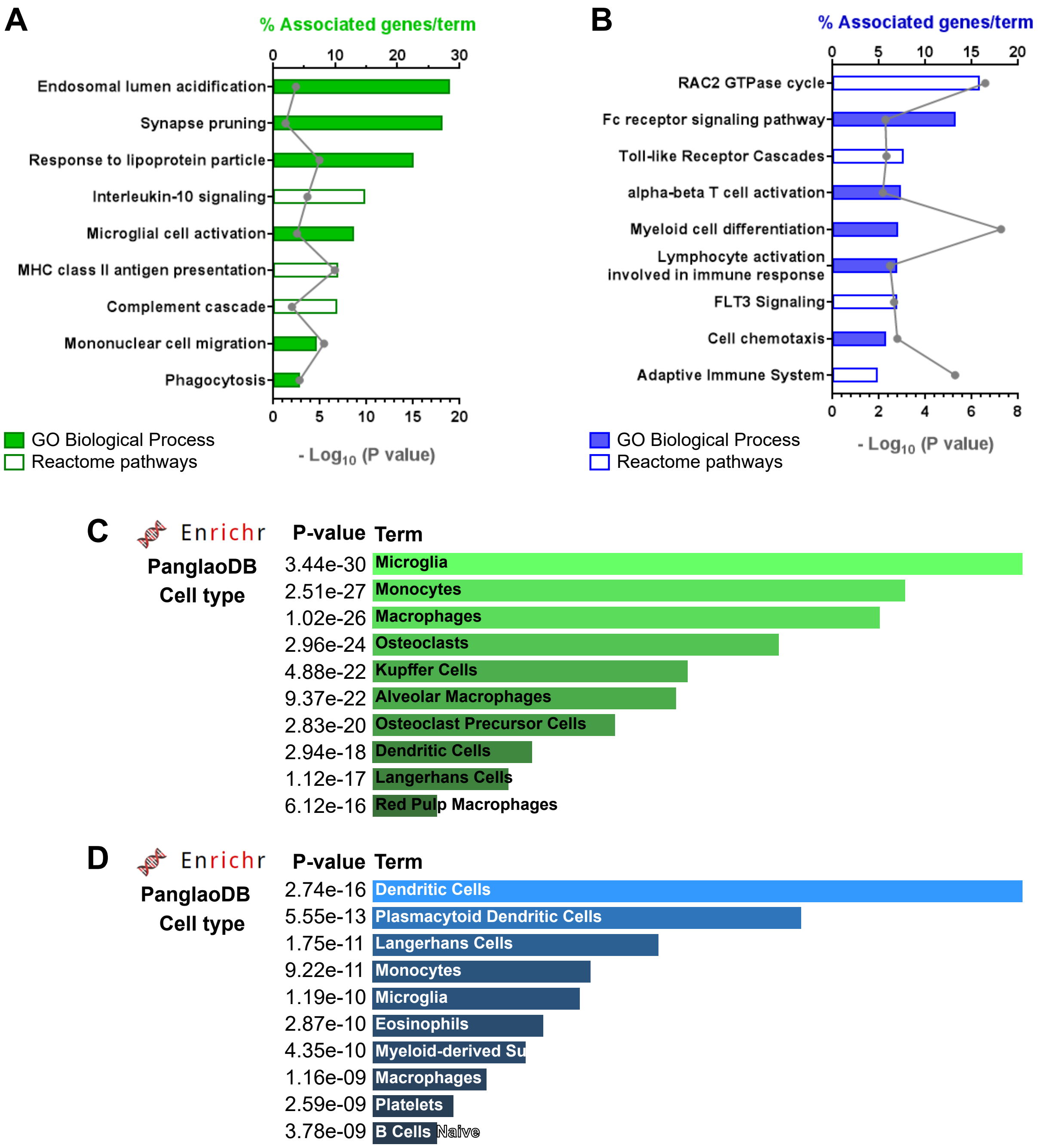


**Figure 4 – figure supplement 2. Functional analysis using the corresponding mammalian orthologs. A-B.** Pathway analysis of the significantly up-regulated markers ($P_{adj}$ <0.05, $log_2fc$=0.25) for microglia (A) and DC-like (B) clusters. The histogram bars represent the percentage of the total number of associated genes per term found and the line graph represents the level of statistical significance ($P_{adj}$ $-log_{10}$) of the enriched GO terms and Reactome pathways. **C-D.** Cell type enrichment analysis using PanglaoDB from the Enrichr engine and uploading the same gene list as in A-B.

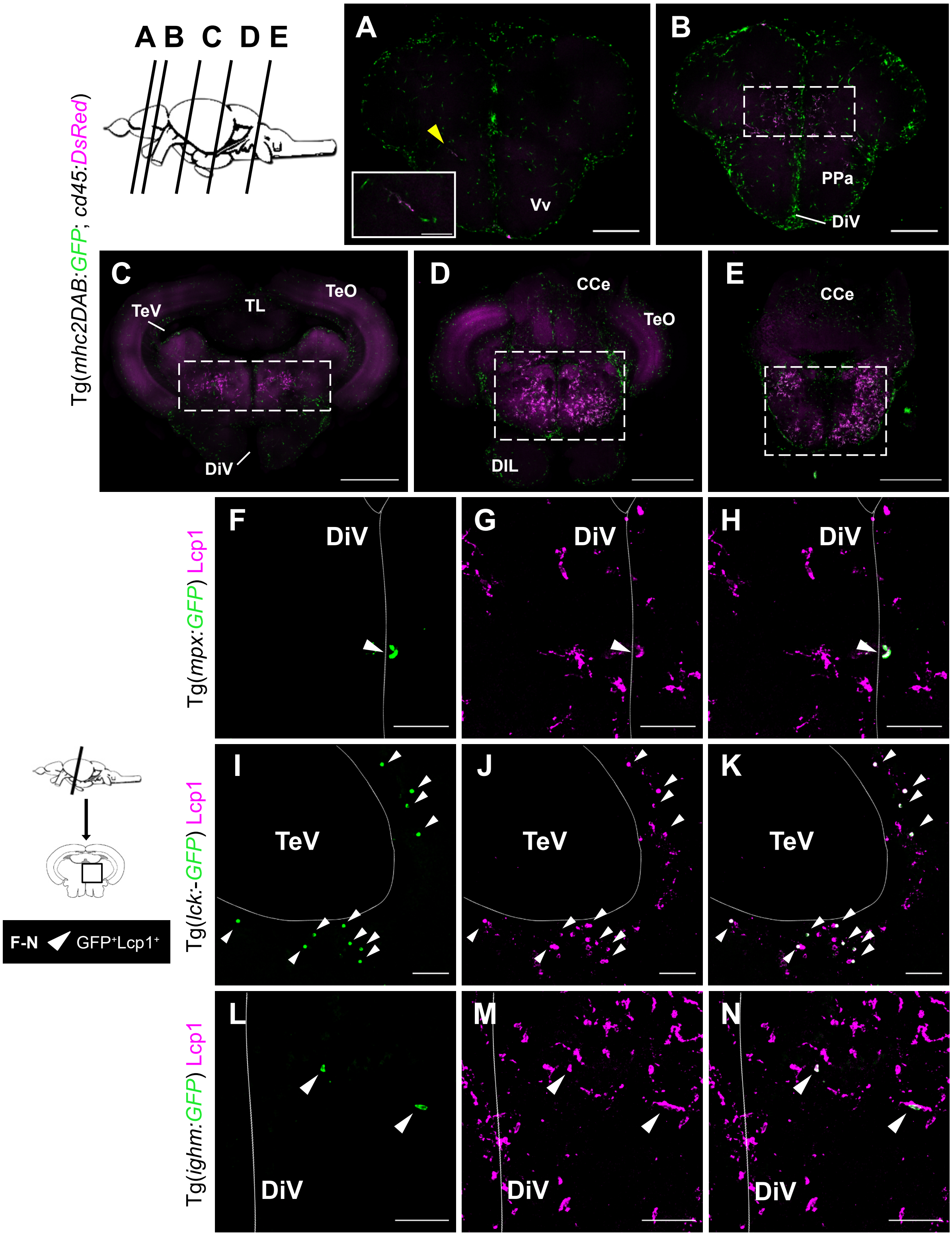


**Figure 5 – supplement 1. Distribution of DC-like cells and immunofluorescence staining for neutrophils and lymphoid cells in the adult brain. A-E.** Vibratome sections (100 µm) from an adult *Tg(mhc2dab:GFP; cd45:DsRed)* brain. **A-B.** Anterior telencephalon sections contain few *cd45$^{high}$; mhc2$^{+}$* cells (yellow arrowhead) and these are more abundant in posterior telencephalic sections (dashed area). Scale bars: 200 µm. **C-D.** Posterior midbrain sections and hindbrain (**E**). Scale bars: 500 µm. DiV, diencephalic ventricle; TeV, telencephalic ventricle; Vv, ventral nucleus of ventral telencephalic area; PPa, parvocellular preoptic nucleus anterior part; TL, torus longitudinallis; OTe, tectum opticum; DIL, lobus caudali cerebelli; CCe corpus cerebelli. **F-N.** Blood lineage-specific reporter lines labeling neutrophils (*Tg(mpx:GFP)*) (F-H), T and NK cells (*Tg(lck:GFP)*) (I-K) and B lymphocytes (*Tg(ighm:GFP)*) (L-N) were used in these experiments. Sections were co-stained for GFP (green, left panels) and the pan-leukocytic marker L-plastin (Lcp1) (magenta, middle panels) to validate the hematopoietic identity of GFP-expressing cells. Merged images are represented in the right panels. Representative midbrain sections are shown. **F-H.** In line with our flow cytometry analyses, neutrophils are scarce in the zebrafish brain. A rare neutrophil is shown lining the borders of the diencephalic ventricle (DiV). **I-K.** T/NK cells are mainly located in the periphery or lining the ventricles (dashed line) and occasionally within the brain parenchyma. **L-N.** B cells are rarely found in the brain or within the brain parenchyma. Scale bars: 50 µm.

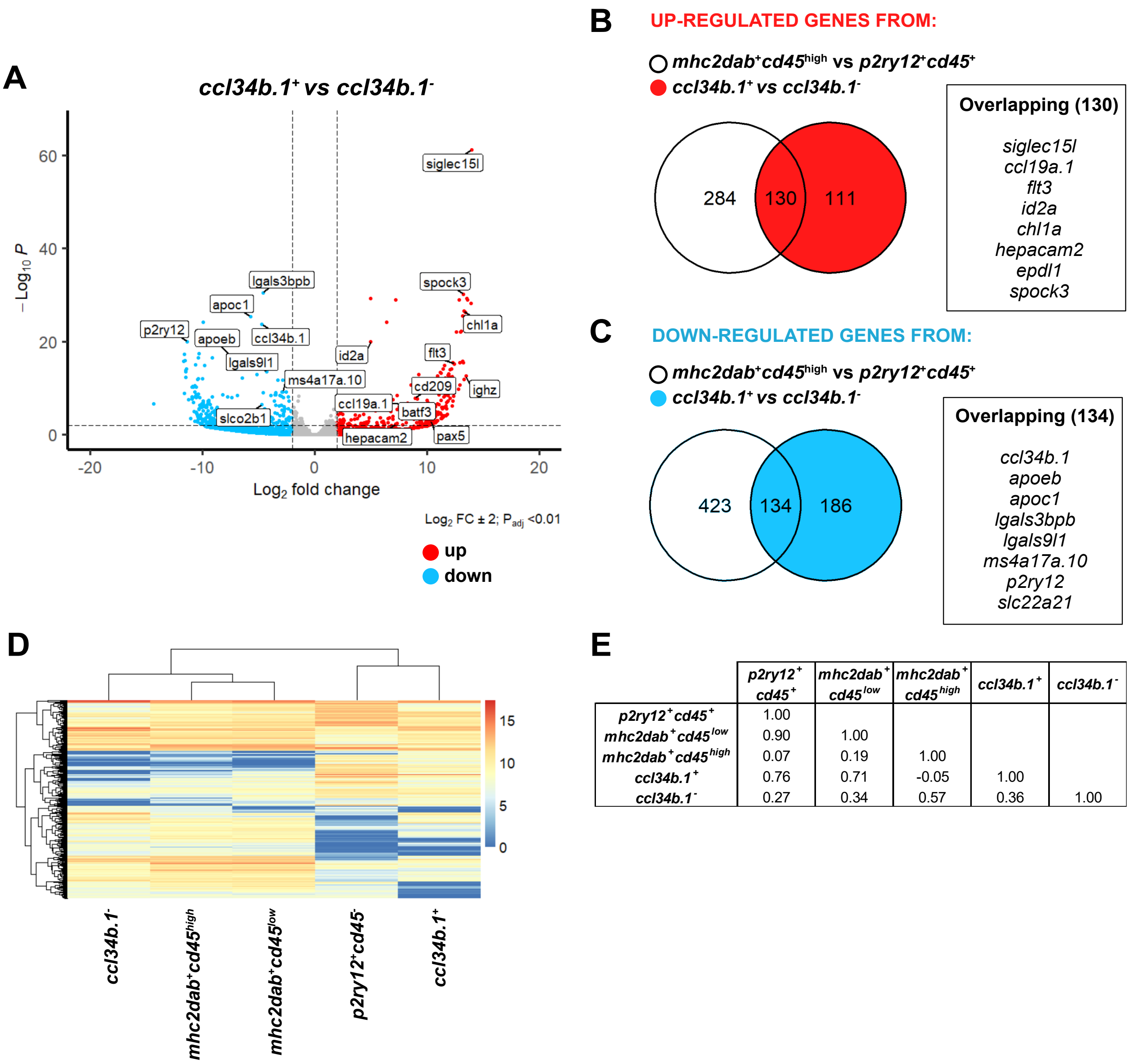


| | $p2ry12^{+}$ $cd45^{+}$ | $mhc2dab^{+}$ $cd45^{low}$ | $mhc2dab^{+}$ $cd45^{high}$ | $ccl34b.1^{+}$ | $ccl34b.1^{-}$ |
|---|---|---|---|---|---|
| ***p2ry12*⁺*cd45*⁺** | 1.00 | | | | |
| ***mhc2dab*⁺*cd45*^low^** | 0.90 | 1.00 | | | |
| ***mhc2dab*⁺*cd45*^high^** | 0.07 | 0.19 | 1.00 | | |
| ***ccl34b.1*⁺** | 0.76 | 0.71 | -0.05 | 1.00 | |
| ***ccl34b.1*⁻** | 0.27 | 0.34 | 0.57 | 0.36 | 1.00 |

**Figure 6 – figure supplement 1. Differential expression analysis reveal that brain *ccl34b.1:GFP*⁻*; mpeg1.1:mCherry*⁺cells have a DC-like transcriptome . A.** Volcano plot showing the DE genes between two parenchymal populations previously described in Wu et al. (2020) as *ccl34b.1*⁺; *mpeg1.1*⁺ phagocytic microglia and *ccl34b.1*⁻; *mpeg1.1*⁺ regulatory microglia. Red dots represent up-regulated genes and blue dots down-regulated genes. Lines indicate significantly DE genes ($log_2$ fold-change >|2|, $-log_{10}$ $P_{adj}$ <0.01). Labels show that these cells differentially express marker genes identified for DC-like and microglia in our scRNA-sequencing analysis. **B.** Venn diagram showing 130 overlapping genes amongst the significantly up-regulated DE genes ($log_2$ fold-change >|1|, $-log_{10}$ $P_{adj}$ <0.01) between DC-like and microglia from our dataset and *ccl34b.1*⁻; *mpeg1.1*⁺ and *ccl34b.1*⁺; *mpeg1.1*⁺ cells from Wu et al (2020). The majority of DC-like marker genes are shared, suggesting that DC-like and *ccl34b.1*⁻*mpeg1.1*⁺ cells have a similar expression profile. **C.** Venn diagram showing 134 overlapping genes amongst the significantly down-regulated DE genes between DC-like cells and microglia from our dataset and *ccl34b.1*⁻; *mpeg1.1*⁺ and *ccl34b.1*⁺; *mpeg1.1*⁺ cells from Wu et al (2020). The majority of microglia marker genes are shared, suggesting that microglia and *ccl34b.1*⁺; *mpeg1.1*⁺ cells have a similar expression profile. **D.** Hierarchical clustering of the normalized expression of the top DE expressed genes in each sample (a total of 1252 genes; $log_2$fc 1.2 $P_{adj}$<0.01). DC-like cells (*mhc2dab*⁺*; cd45*^high^) cluster together with *ccl34b.1*⁻; *mpeg1.1*⁺ cells, while microglia (either as *p2ry12*⁺*; cd45*⁺ or *mhc2dab*⁺*; cd45*^low^) cluster together with *ccl34b.1*⁺; *mpeg1.1*⁺ cells. Color scale indicates normalized expression for each gene. **E.** Correlation matrix (spearman) performed using the list of the top DE expressed genes (a total of 1252 genes; $log_2$fc 1.2 $P_{adj}$ <0.01) shows that DC-like and *ccl34b.1*⁻; *mpeg1.1*⁺ cells are positively correlated, suggesting their similar identity, while they show a decreased relationship with microglia (*p2ry12*⁺*; cd45*⁺ or *mhc2dab*⁺*; cd45*^low^) and *ccl34b.1*⁺; *mpeg1.1*⁺ cells.

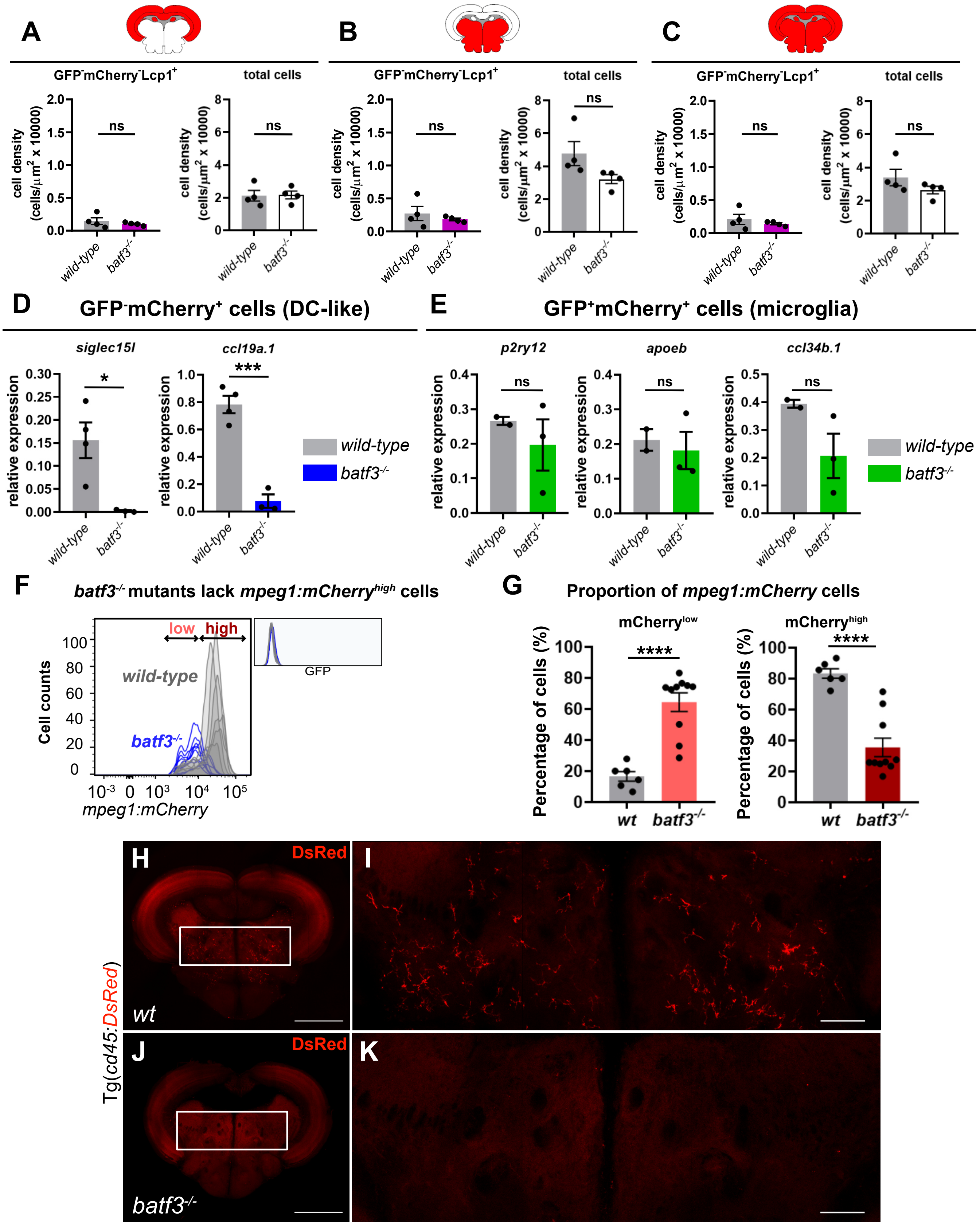


**Figure 7 – figure supplement 1. Characterization of brain immune cells in the *batf3*$^{-/-}$ mutant. A-C.** Cell density quantification of non myeloid cells (GFP$^-$; mCherry$^-$; Lcp1$^+$) and total leukocyte(all Lcp1$^+$) in the dorsal midbrain area or optic tectum (A), ventral midbrain area (B) and the whole section (C) of *wild-type* and *batf3*$^{-/-}$ fish from Figure 7. Data represent mean ± SEM (*n*=4 fish). **D,E.** Q-PCR expression for DC-like markers *siglec15l* and *ccl19a.1* in *p2ry12:GFP*$^-$*; mpeg1:mCherry*$^+$ DC-like cells (D) and for microglia-specific genes *p2ry12*, *apoeb* and *ccl34b.1* in *p2ry12:GFP*$^+$*; mpeg1:mCherry*$^+$ microglia (E) sorted from *wild-type* (grey bars) and *batf3*$^{-/-}$ (colored bars) adult brains. * P<0.05, *** P<0.001 (Two-tailed unpaired t-test). Data are represented as mean ± SEM. *n* refers to the number of biological replicates. **F.** Number of cells (y-axis) versus mCherry fluorescence intensity (x-axis) histogram plot of brain cell suspensions from controls (grey) and *batf3*$^{-/-}$ (blue) fish carrying the *Tg(p2ry12:GFP; mpeg1:mCherry)* double transgene. It shows that all residual *mpeg1:mCherry*$^+$ cells in the *batf3* mutant display lower fluorescence intensity as compared to their *wild-type* counterparts. **G.** Proportion of *mpeg1:mCherry*$^+$ cells according to their low or high fluorescence intensity as shown in F. Data represent mean ± SEM (*n*=4). *n* refers to the number of biological replicates. **** P<0.0001 (Two-tailed unpaired t-test). **H-K.** Illustrative case of vibratome midbrain sections of *Tg(cd45:DsRed)* transgenic *wild-type* (H,I) and *batf3*$^{-/-}$ (J,K) fish (*n*=3). The endogenous fluorescence of DsRed$^{high}$ DC-like cells permits their identification in the ventral part of *wild-type* brains (*n*=3). However, these cells are not found in the *batf3*$^{-/-}$ mutant (*n*=3). Scale bar: 500 µm. **I, K.** High magnification of the insets (white frame) in H (I) and J (K). Scale bar: 100 µm. Note that cells with lower fluorescence intensity (e.g. cd45low microglia) are not detectable using this approach due to signal loss (quenching) following fixation.

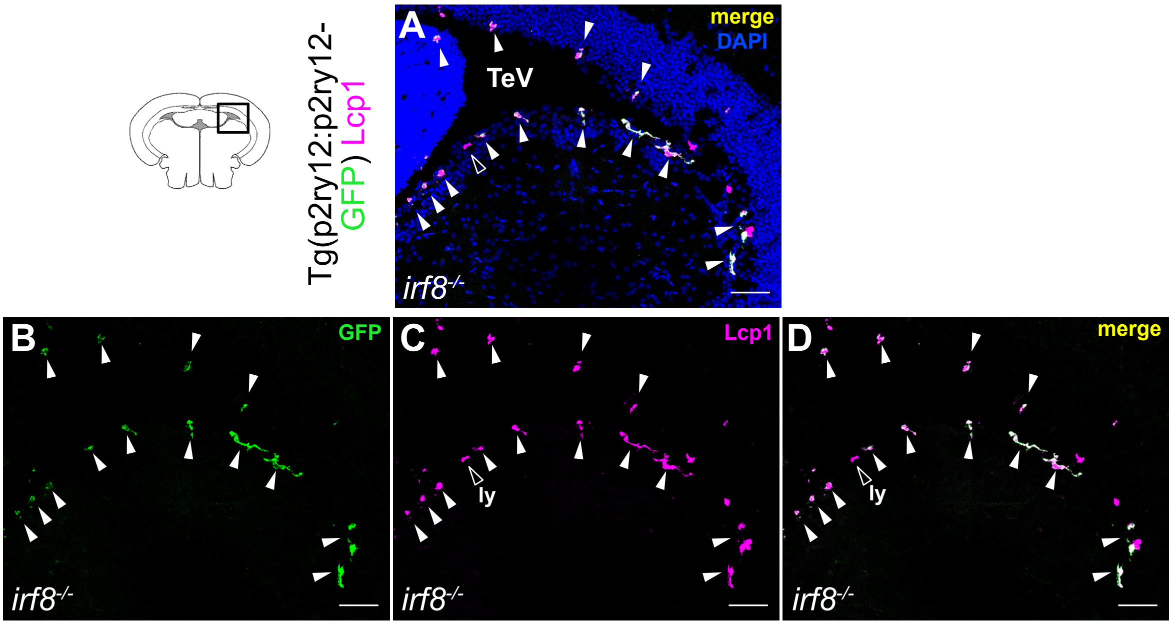


**Figure 8 – supplement 1. Location of microglia in the brain ventricles of *irf8*-deficient fish. A-D.** Immunofluorescence of brain sections from *irf8*$^{-/-}$ mutant fish carrying the *Tg(p2ry12:p2ry12-GFP)* reporter, co-stained with GFP (green) and Lcp1 (magenta) antibodies. **B.** Microglial cells in the mutant are mostly found lining the ventricles (A-D, white arrowheads). Nuclear DAPI staining is shown in A to better visualize the position of the GFP$^+$ and Lcp1$^+$ cells along the ventricle. Ly, lymphocyte (outline white arrowhead). Scale bar represents 50 µm. **B.** GFP$^+$ cells. **C.** Lcp1$^+$ cells. **D.** merged channels. Scale bar: 50 µm. TeV, telencephalic ventricle.

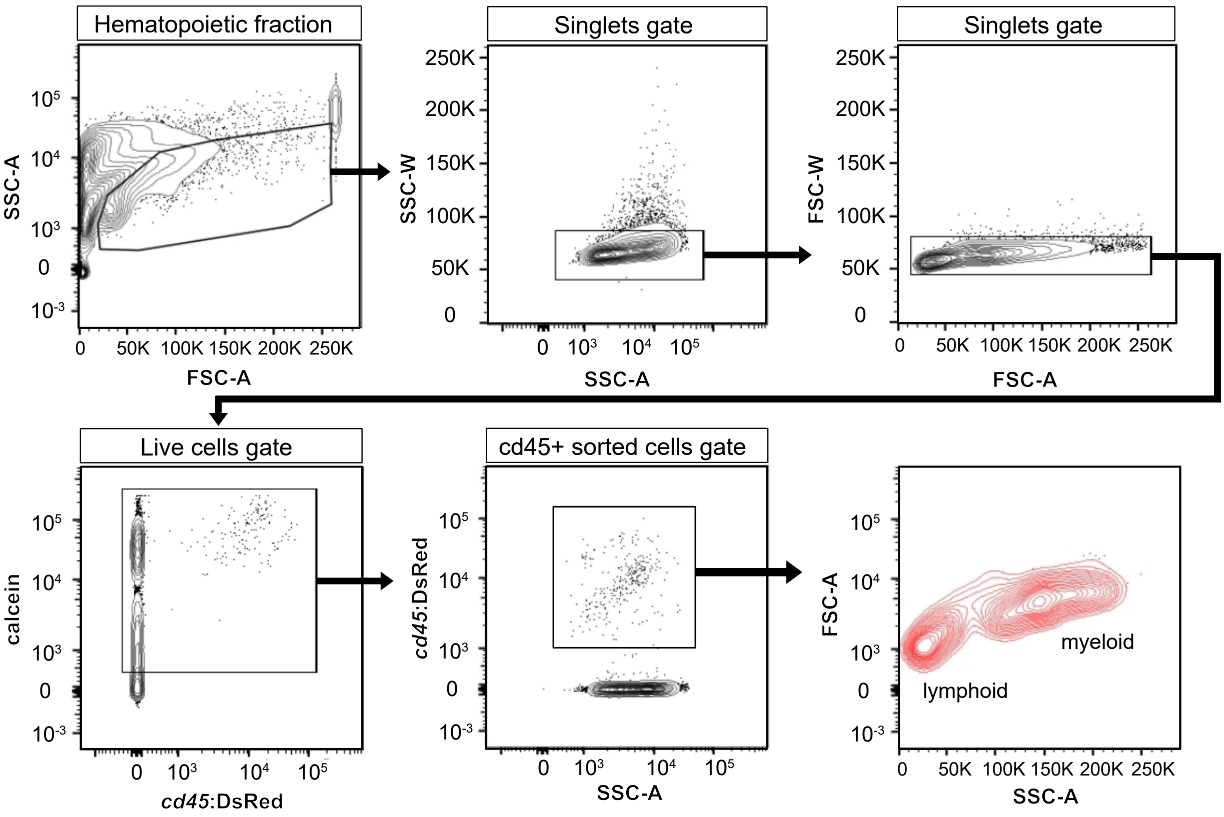


**Appendix 1 – Figure 2. Isolation of a pure population of brain leukocytes.** FACS plots illustrating the gating strategy to isolate total brain leukocytes from *Tg(cd45:DsRed)* transgenic fish. Cells are first selected based on side (SSC-A) and forward scattering (FSC-A) characteristics. Cell doublets are then filtered out based on the width of the side scatter and forward signal. Next, calcein$^+$ live cells are selected and *cd45:DsRed*$^+$ cells are sort out of the live population. As shown in the last plot, this strategy captures leukocytes from both the lymphoid and myeloid lineages, with the exception of B cells, as they don't express the *cd45:DsRed* transgene. In these experiments, brain *cd45:DsRed*$^+$ cells isolated from 3 individual fish were pooled for scRNA sequencing.

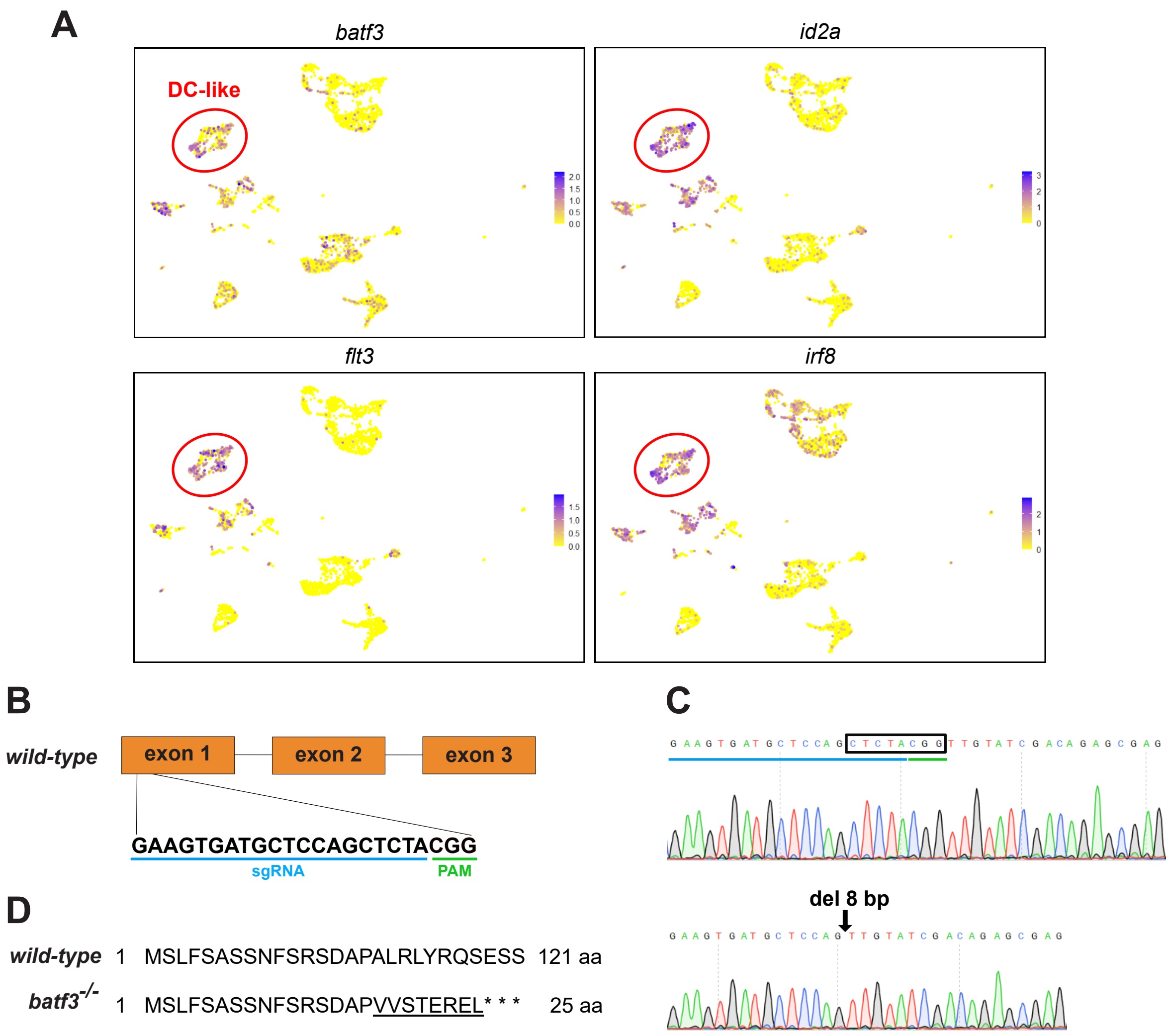


**Appendix 2 - Figure 7. Generation of *batf3*$^{-/-}$ CRISPR mutants. A.** UMAP visualization of the expression pattern of selected zebrafish DC-like cluster genes whose orthologs represent known markers of cDC1 in mammals, including *batf3*, *id2a*, *flt3* and *irf8*. Color scale (from yellow to purple) indicates the expression level for each gene (normalized counts in log1p). **B.** Schematic view of exons 1 to 3 of the *batf3* gene, with the CRISPR/Cas9 targeting sequence in exon 1 underlined in blue and the PAM sequence underlined in green. **C**. Sequence chromatograms of *wild-type* (upper row) and mutant (lower row) sequences, showing the 8-bp deletion (black box) in *batf3*$^{ulb31}$ fish. **D.** Predicted Batf3 protein sequence in the *wild-type* and the mutant. The 8-bp deletion in the *batf3* gene induces a frameshift (underlined) that results in the production of a truncated protein due to the presence of 3 consecutive premature stop codons.
